# Mathematical model shows how sleep may affect amyloid *β* fibrillization

**DOI:** 10.1101/751230

**Authors:** Masoud Hoore, Sahamoddin Khailaie, Ghazal Montaseri, Tanmay Mitra, Michael Meyer-Hermann

## Abstract

Deposition of amyloid β (Aβ) fibers in extra-cellular matrix of the brain is a ubiquitous feature associated with several neurodegenerative disorders, especially Alzheimer’s disease (AD). While many of the biological aspects that contribute to the formation of Aβ plaques are well addressed at the intra- and inter-cellular level in short timescales, an understanding of how Aβ fibrillization usually starts to dominate at a longer timescale in spite of the presence of mechanisms dedicated to Aβ clearance, is still lacking. Furthermore, no existing mathematical model integrates the impact of diurnal neural activity as emanated from circadian regulation to predict disease progression due to a disruption in sleep-wake cycle. In this study, we develop a minimal model of Aβ fibrillization to investigate the onset of AD over a long time-scale. Our results suggest that the diseased state is a manifestation of a phase change of the system from soluble Aβ (sAβ) to fibrillar Aβ (fAβ) domination upon surpassing a threshold in the production rate of soluble Aβ. By incorporating the circadian rhythm into our model, we reveal that fAβ accumulation is crucially dependent on the regulation of sleep-wake cycle, thereby indicating the importance of a good sleep hygiene in averting AD onset. We also discuss potential intervention schemes to reduce fAβ accumulation in the brain by modification of the critical sAβ production rate.

## Introduction

Amyloid *β* (A*β*) accumulation is a common pathological characteristic of several neurodegenerative and neuroinflammatory disorders, essentially Alzheimer’s disease (AD) [1, 2]. The clinical symptoms and cellular dysfunction that contribute towards pathological burden in AD are generally considered to be the outcome of neurodegenerative effects of the formation of A*β* plaques in the extra-cellular space [3] and neurofibrillary tangles of *τ* protein in neurons [4]. Regulation of A*β* levels in the interstitial fluid (ISF) is mediated by circadian rhythm [5], which also modulates the neuro-immune-endocrine system through several complex biochemical mechanisms [6, 7, 8]. Thus, it might be possible that the circadian rhythm and the emergence of neurodegenerative disorders have a bidirectional modulatory relationship with each other. For example, the sleep-wake cycle is known to be altered several years before the onset of AD [9, 10], and prolonged neuroinflammation may also contribute to worsen the sleep hygiene.

It is often debated whether alteration in the sleep rhythm is a causal player in AD, or it only correlates with it [11, 9, 10]. There exists a positive correlation between the circadian gene expression and the number of hypothalamic neurons in the suprachiasmatic nucleus (SCN) that acts as a master regulator of circadian rhythm [12, 13]. An intriguing fact in this regard is that a significant loss of neurons is found in the SCN of the mammals debilitated with AD [14], which may suggest why the sleep-wake cycle is disrupted in a broad range of neuroinflammatory disorders. Moreover, a disturbance in the sleep hygiene is found to correlate with A*β* deposition in the brain [15].

A*β* is produced by neurons upon synaptic activity and released into the ISF [16, 17], thereby having an interplay with neural activity. As the SCN controls diurnal neural activity through circadian rhythm, the secretion of A*β* into the ISF follows a similar diurnal rhythm in normal circumstances. This implies that a disturbance in the sleep-wake cycle or unchecked deprivation in sleep may result in higher A*β* production through higher neural activity and may also lead to A*β* neurotoxicity and oxidative stress to neurons. As a consequence, a disrupted central circadian rhythm results in altered hippocampal A*β* rhythm and causes accumulation of amyloid plaques. A*β* plaques that contribute to neurotoxicity disturb the neural function in the SCN and impair the regulation of circadian neural activity throughout the brain [12]. This acts as a positive-feedback loop resulting in more A*β* accumulation in the SCN, more neurotoxicity to the SCN, less regulation of the global brain diurnal rhythm, and A*β* secretion as well as subsequent fibrillization.

As there are a plethora of factors that can contribute to neuroinflammation and neurodegeneration as we age, it is practically impossible to take all the details in shorter timescales (e.g. milliseconds in case of neuronal firing rate), and draw a broad unified picture of the disease development over an extremely long timescale, i.e., in years. Thus, it will be intriguing to design a mean-field approach where one can, starting from the key components of the underlying system, understand the transition into the diseased state characterized by A*β* fibrillization long before the patho-physiological symptoms occur and neuroinflammatory response starts. We propose a minimal model and show that the onset of AD is driven by a phase change from the soluble form of A*β* to its fibrillar form. The proposed model has the advantage of simplicity compared with the recent mathematical models [18, 19, 20, 21]. A few other in-silico studies explained how the A*β* fibres form in the brain, using an agent-based model [22] or numerically solving the partial differential equations [21, 23]. Although many aspects of the disease are explained and understood from these models, we still lack a simple explanation of how different factors affect the brain in the course of AD.

In this paper, we employ a minimal mathematical model based on ordinary differential equations (ODEs) to investigate A*β* fibrillization in the brain, and its dependence on associated factors such as A*β* production by neurons and their clearance by microglia or efflux through the cerebrospinal fluid (CSF). Our results suggest that the accumulation of fibrillar A*β* can be viewed as a shift in the scaling law between the concentration of fibrillar A*β* and soluble A*β* production rate due to neuronal activity. Our work also predicts that even a two-fold increase in A*β* production may trigger a phase change from soluble A*β* to fibrillar A*β*. The current ODE-based model suggests that AD can be seen as the outcome of perturbations in brain homeostasis. We believe that such a minimal model could also pave a way for new potential treatments or prevention strategies.

## Methods

We base the mathematical model on major mechanisms regulating the fibrillization process, namely fibrillar A*β* formation from soluble A*β*, A*β* production by neurons, and their clearance by glial cells or efflux through the CSF.

The interaction network of the fibrillization of A*β* in AD is depicted in Fig. 1. The changes in the number and activity of astrocytes, neurons, and microglia are ignored. The cycling time of microglia is much faster (a few days) than the time scale of A*β* accumulation (a few years). Thus, we assume that the number of activated microglia reaches its carrying capacity through a fast logistic growth due to proliferation in the presence of sA*β* and fA*β*. We assume that the number of neurons do not vary significantly over the course of the onset of A*β* fibrillization [38, 39] such that the production rate of A*β*, *R*, remains stable for a long period of time. With the aforementioned biological assumptions, the equation for the rates of change in A*β* concentrations in the ODE-based model, depicted in Fig. 1, can be obtained as

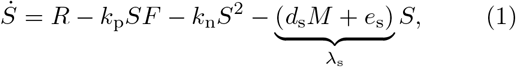

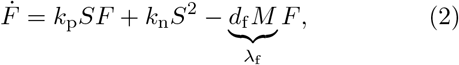

where *S* and *F* are the concentration of A*β* oligomers in soluble (*S ≡* [sA*β*]) and fibrillar form ([*F ≡* [fA*β*]]), respectively. *λ*_s_ = *d*_s_*M* +*e*_s_ and *λ*_f_ = *d*_f_ *M*, where *M* represents the number density of microglia. Over-dots denote time derivative (*Ẋ* d*X/*d*t*). *R* is the rate of sA*β* production over a day due to neural synaptic activity. Although the production rate has a circadian pattern, its day average is used as an approximation. *k*_n_ and *k*_p_ are the A*β* nucleation and polymerization rates, respectively. The parameter values are provided in Tab. 1.

**Figure 1:**
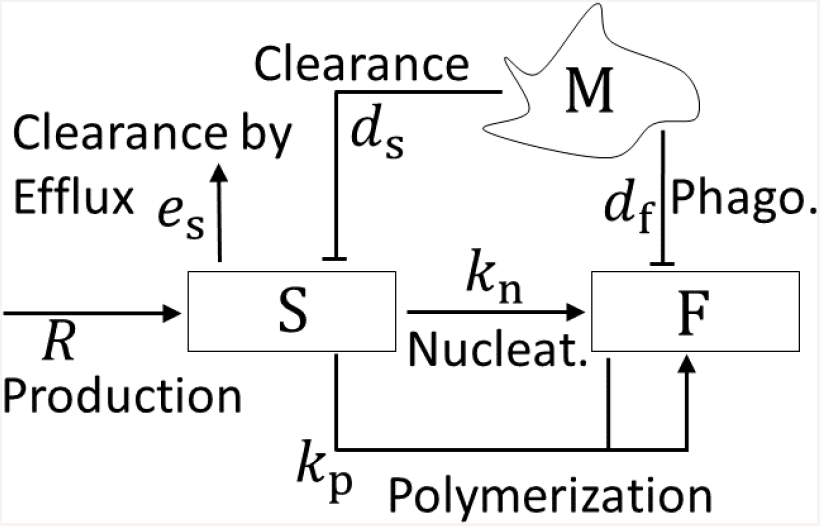
Model. Signaling network of A*β* fibrillization. sA *β* (*S*) is produced by the mean rate *R*, and cleared by efflux through the CSF. *S* transforms to fA*β* (*F*) by nucleation and polymerization. Microglia (*M*) clear *S* and *F* through macropinocytosis and phagocytosis, respectively.

**Table 1:** Model parameters and their estimated values. It is noted that for all the analysis in this work, the parameters are taken from this table unless otherwise specified.

| Symbol | Description | Estimated value | Unit | Ref. |
| --- | --- | --- | --- | --- |
| $S$ | concentration of $A\beta$ oligomers in soluble form | variable | $\mu\text{M}$ | |
| $F$ | concentration of $A\beta$ oligomers in fiber form | variable | $\mu\text{M}$ | |
| $R$ | mean rate of $sA\beta$ secretion by neurons | variable | $\mu\text{M} \cdot \text{day}^{-1}$ | [24] |
| $a$ | normalized amplitude of circadian rhythm in $sA\beta$ production | 25% | - | [25, 24, 26, 27] |
| $\tau$ | period of circadian rhythm | $\sim 1$ | day | |
| $M$ | number density of microglia, $N_{\text{mgl}}/V_{\text{brain}}$, assuming that microglia/neuron ratio is unity | $\sim 6.7 \times 10^7$ | $\text{mL}^{-1}$ | [28, 29] |
| $k_p$ * | rate of $A\beta$ fiber growth | $2.6 \times 10^1$ | $\mu\text{M}^{-1} \cdot \text{day}^{-1}$ | [30] |
| $k_n$ * | rate of $A\beta$ fiber nucleation | $1.7 \times 10^{-1}$ | $\mu\text{M}^{-1} \cdot \text{day}^{-1}$ | [30] |
| $d_s$ | rate of $sA\beta$ clearance by microglia | $5.3 \times 10^{-9}$ | $\text{day}^{-1} \cdot \text{mL}$ | [31] and [32, 33] |
| $e_s$ | rate of $sA\beta$ efflux through the CSF | 1.9 | $\text{day}^{-1}$ | [34, 35] |
| $d_f$ | clearance of $fA\beta$ by microglia | $\sim d_s/30$ | | [36] |

\* Estimation of $k_n$ and $k_p$ has been done using the differential evolution method [37] (see Fig. S1).

The biological observations and associated modeling assumptions are explained as follows.

### sA*β* efflux

It has been found that meningeal lymphatics plays an important role in the clearance of the brain waste including sA*β* [40, 41, 42]. The efflux of sA*β* to the meningeal lymphatics and CSF is independent of the existence of microglia and is driven by diffusion. The efflux rate of sA*β* in the CSF, *e*_s_, has been reported to be around 7.6 to 8.3 % per hour [34, 35] (i.e. *e*_s_ = 1.9 *±* 0.4 day^*−*1^).

### A *β* clearance by microglia

Fibrillar A*β* (fA*β*) is cleared by microglia through phagocytosis while its soluble form (sA*β*) is cleared through macropinocytosis [43, 31], a process mediated by cellular internalization. Macropinocytosis is linearly dependent on the concentration of the entities that are internalized by the cell. Thus, the rate of sA*β* clearance does not saturate because of such a mechanism. In contrast, fA*β* ligates a receptor on microglia and its phagocytosis is better represented by a saturable rate obeying the Michaelis-Menten kinetics [44]. Nevertheless, the clearance or decay rates of sA*β* and fA*β* are modeled similarly, assuming that the number of microglia receptors is much greater than the number of A*β*, while keeping in mind that such an assumption is not valid at high concentrations of fA*β*. However, it applies well to the early stage of the disease. The concentration of sA*β*, *S ≡* [sA*β*] and fA*β*, *F ≡* [fA*β*] decrease in time by the rates *d*_s_*M* and *d*_f_ *M*.

It was observed that the intracellular concentration of sA*β* depends linearly on the concentration of sA*β* in the tissue (as a result of macropinocytosis) [31]. Furthermore, the intracellular concentration of sA*β* decays exponentially. With this, the clearance rate of sA*β* by microglia is estimated as 0.22 h^*−*1^*N*_mgl_*V*_mgl_*/V*_brain_, assuming a 100% uptake by microglia, with *N*_mgl_, *V*_mgl_ and *V*_brain_ being the total number of microglial cells, the volume of microglial cell and the total volume of the brain, respectively. Microglia have a plethora of shapes and sizes according to their activation states [32, 33] making the estimation for *V*_mgl_ challenging. Here, it is assumed that *V*_mgl_ is in the order of 1000 *μ*m^3^, leading to *d*_s_ = 0.22 *×* 10^*−*9^ h^*−*1^mL. It is roughly estimated that *d*_f_ is 30 times smaller than *d*_s_ [36]. With the estimated values, microglial clearance composes 20% of the whole clearance, while the efflux has a greater share of 80%.

### Fibrillization process

Shortly after the introduction of amyloid cascade hypothesis [45], the fibrillization of A*β* has been investigated extensively by many scientists [46]. According to the seminal model for amyloid fibrillization proposed by Lomakin et al. in 1997 [47], the accumulation of A*β* is driven through two distinct pathways, the initial nucleation and the secondary nucleation by adjoining of soluble A*β* to amyloid fibers [47]. Although the amyloid aggregation is still an active field of research, there is a consensus that such a phenomenon occurs through these two mechanisms. The secondary nucleation has been understood as the leading mechanism for amyloid plaque formation in AD [48, 49, 50].

We take the most simple form of amyloid aggregation, considering a primary and secondary nucleation, referred to as nucleation and polymerization, respectively. In the primary nucleation, soluble A*β* come together and form small fibers with the nucleation rate *k*_n_. In the secondary nucleation, soluble A*β* adjoin the fibers with polymerization rate *k*_p_. The required energy for A*β* nucleation is one order of magnitude higher than for their polymerization [51], implying that the polymerization rate is much greater than the nucleation rate (*k*_p_ ≫ *k*_n_). In order to calibrate the model parameters with the experimental results, the recent ThT experiments of amyloid aggregations [30], upon which many kinetic models of polymer aggregation are being validated [52, 53, 54] have been used. The estimated values for *k*_n_ and *k*_p_ are tabulated in Tab. 1 (see also Fig. S1).

The large difference between the values of *k*_p_ and *k*_n_, and the scaling of A*β* nucleation with *S*^2^ due to their self-interaction are essential in determining an abrupt phase change from soluble to fibrillar A*β*. Such a fibrillization dynamics is in qualitative agreement with the complex kinetic models for amyloid fibrillization [55, 54, 53, 56, 47].

## Results and Discussion

### The emergence of a soluble and a fibrillar phase with distinct scaling laws

We assume that a healthy brain in its initial state does not have any fA*β* and is characterized by sA*β* production due to neuronal activity. Formation of sA*β* by neurons sets A*β* dynamics in the system and initiates the interplay between several biological processes as discussed above. As the “diseased” state of the brain in AD is marked by a high concentration of fA*β*, it becomes primarily important to understand how the dynamics of the system may result in fA*β* formation in the extracellular matrix and how it would depend on the production rate of sA*β*, *R*. To perceive this, we first calculate the steady state of our model system and depict its dependence on *R* (Fig. 2). Considering that the system starts with zero concentration of fA*β*, the steady-state solutions (*S_∞_* ≡ [sA*β*]_*t*→∞_ − *S*_c_ and *F_∞_* ≡ [fA*β*]*t*) of Eq. 1 and 2 read

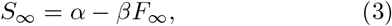

and

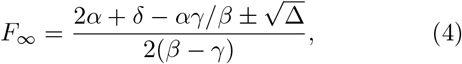

where *α* = *R/λ*_s_, *β* = *λ*_f_ */λ*_s_, *γ* = *k*_p_*/k*_n_, *δ* = *λ*_s_*/k*_n_, and Δ = (*αγ/β*)^2^ + *δ*^2^ + 4*αδ −* 2*δαγ/β*.

**Figure 2:**
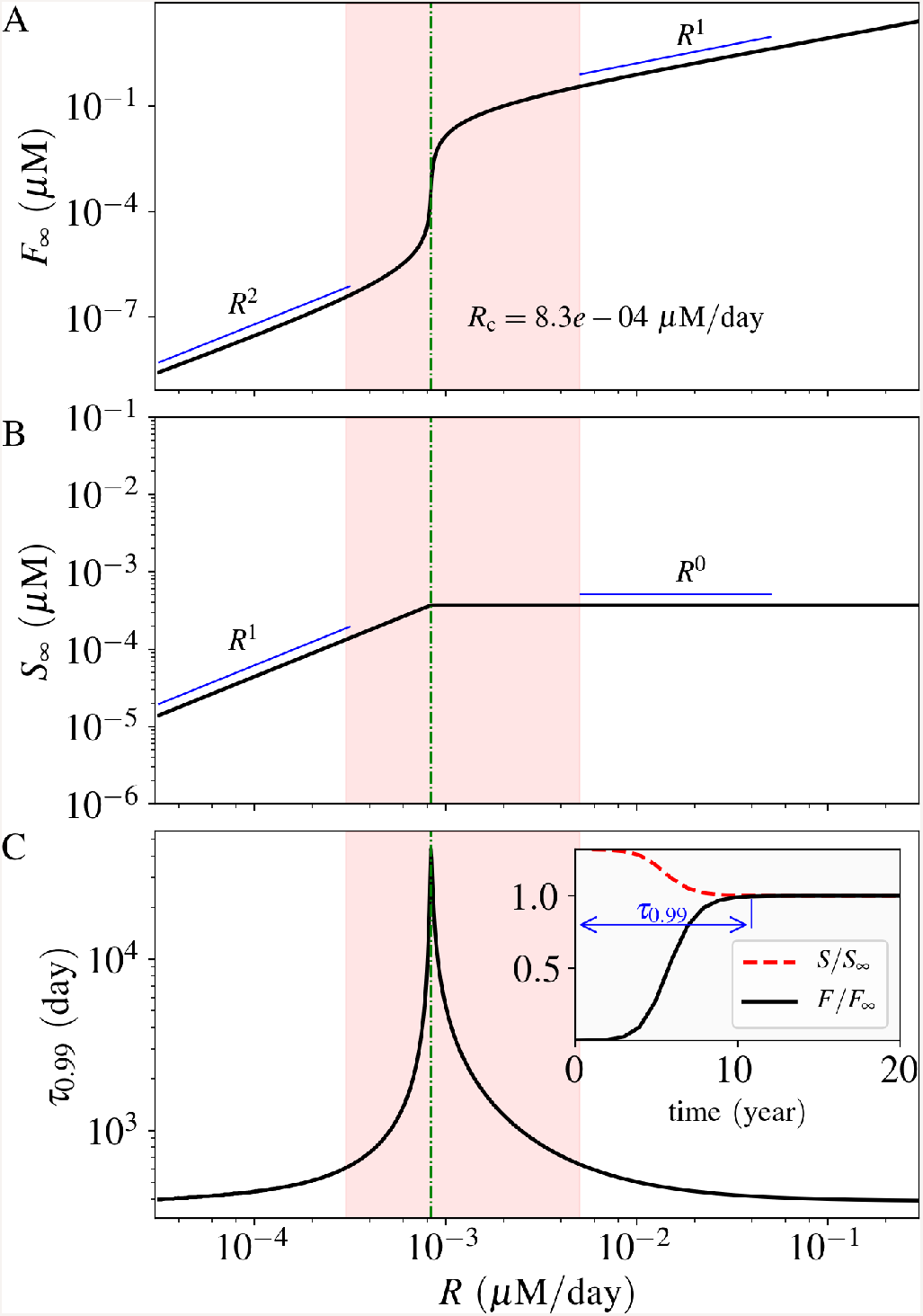
Steady-state solution and settling time. Panels A and B show the steady-state solutions for the concentration of fA *β* and sA *β*, respectively. The critical A *β* production rate *R*c is shown in the diagram. Panel C shows the corresponding settling time, as defined in the inset. It increases abruptly as the production rate gets closer to the critical rate *R*c. The inset diagram shows the normalized concentrations in time for the case *R* = 1.1 *×* 10^*−*3^ *μ*M *·* day^*−*1^. The estimated physiological production rate is shown by a red-highlighted vertical band in each diagram.

We note that Eq. (4) has only one positive solution for the biologically plausible parameter values listed in Tab. 1. The steady-state solutions of the system variables, *F_∞_* and *S_∞_*, are depicted in Fig. 2. We observe that there is a critical A*β* production rate (*R*_c_) at which the scaling relation of these steady-state solutions with respect to *R* changes (see Fig. 2-A for *F_∞_* and Fig. 2-B for *S_∞_*). Mathematically, *R*_c_ is the point where Δ takes its minimum value with respect to *α* (*R*_c_ = *λ*_s_*δβ*(*γ* − 2*β*)*/γ*^2^). Below *R*_c_, *F*_*∞*_ scales as the second power of *R*, reflecting that nucleation is the leading mechanism of fibrillization. Above *R*_c_, the scaling of *F_∞_* changes to the first power of *R*, implying that polymerization (i.e. adjoining of single oligomers to amyloid fibers) is the leading mechanism. *S_∞_* scales with respect to *R* with an exponent of unity below *R*_c_ and zero above it. It means that the system of ODEs predicts a saturation point for sA*β*. The system has two distinct phases, a phase dominated by soluble A*β*, and another dominated by A*β* in fiber form. As depicted in Fig. 2-A, the fA*β* concentration between the two phases differs by many orders of magnitude, which suggests a clear distinction of the “diseased” state as compared to the “healthy” one.

The time that the system variable *F* takes to reach 99% of its steady-state value is defined as the settling time *τ*_0.99_, as depicted in the inset diagram of Fig. 2-C. The settling time is a measure of how fast the system reaches its steady-state.

We note that the shortest settling time is around one year while it exceeds 100 years as the system moves toward the critical value of the A*β* production rate *R*_c_ (see Fig. 2-C). As can be seen from Fig. 2-C, *τ*_0.99_ varies sharply at the close proximity to *R*_c_, whereas it retains lower values in either side of *R*_c_ and away from it.

We next estimate the biologically plausible range of *R* in the case of humans. It has been shown that A*β* levels in the human brain fluctuate with a diurnal rhythm, whose mesors are around 2.3 × 10^*−*4^ *μ*M for A*β*42 and 2.6 × 10^*−*3^ *μ*M for A*β*40, with normalized amplitudes of 10 to 20% [24]. The production rate of sA*β* has been reported to be around 6-8% per hour in the CSF [34, 35], implying the physiological plausible range of *R ≈* 3 *×* 10^*−*4^ − 5 *×* 10^*−*3^ *μ*M*/*day (shown by the red-highlighted vertical band in Fig. 2). Please note that *R*_c_ belongs to this range based on the values tabulated in Tab. 1.

### Transformation into fibrillar phase depends critically on the system parameters

As the alteration of the phase of A*β* from soluble to fibrillar crucially depends on the critical sA*β* production rate *R*_c_, it is intriguing to explore the various means to shift the value of *R*_c_ to unravel the impact of the distinct underlying biological processes that give rise to the fibrillar phase dominated by fA*β*. It may also uncover potential medical intervention strategies for the prevention or delay of AD onset. To elucidate this, we have studied how a modification in each of the system parameters, listed in Tab. 1 affects *R*_c_.

As expected, *R*_c_ is dependent on the parameters of the model. It scales inversely with *k*_p_ (Fig. S3), and linearly with *e*_s_ (Fig. S4) and *d*_f_ (Fig. S2). Here, as an example, we investigate the effect of parameter *M* in more details. We observe that *R*_c_ scales superlinearly with *M* as depicted in the inset of Fig. 3-A. The steady-state solutions of the system variables (*F_∞_* and *S_∞_*) are also shown for several values of *M* in Fig. 3. Whereas *F_∞_* decreases as we increase *M* (Fig. 3-A), *S_∞_* shows a complex trend (Fig. 3-B). For a characteristic *R* which is beyond *R*_c_’s for all the microglial densities in consideration, *S_∞_* attains a larger value for a higher microglial density (Fig. 3-B). Furthermore, as can be seen from Fig. 3-C, the settling time (*τ*_0.99_) decreases as *R*_c_ (or *M*) increases. In general, the results suggest that an efficient clearance of sA*β* by microglia and an enhanced sA*β* efflux through CSF diminish the risk of developing AD, whereas a high rate of A*β* fiber growth or higher rate of sA*β* production makes an individual prone to AD.

**Figure 3:**
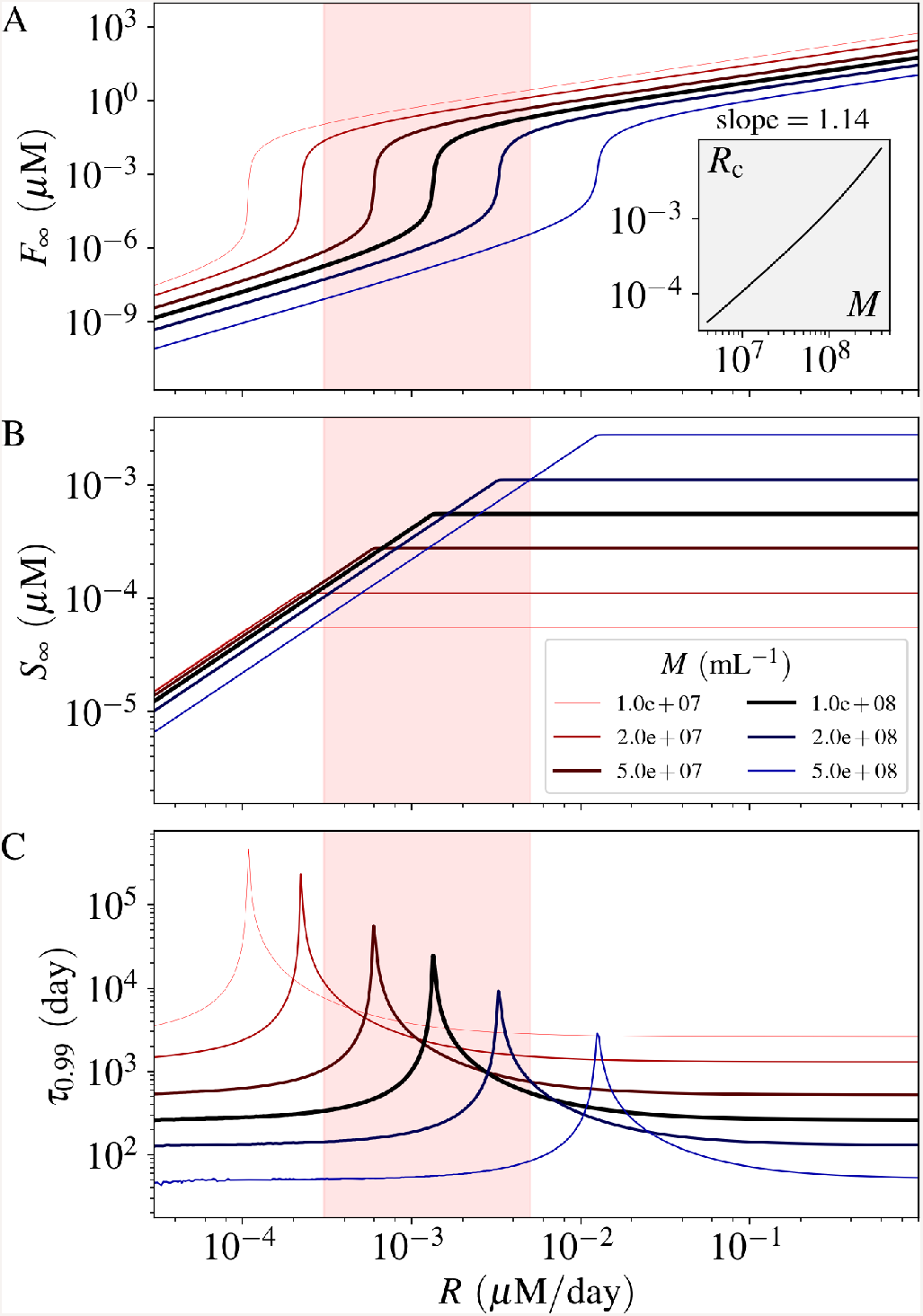
Effect of the number of microglia (*M*) on the fibrillization of A *β*. Similar to Fig. 2, panel A and B show the concentrations and panel C shows the settling time. The inset diagram shows the relation between the critical production rate *R*c and the microglial cell density *M*. The estimated physiological production rate is shown by a red-highlighted vertical band in each diagram. See Fig. S2, S3, and S4 for the impact of other parameters.

### Time-dependent parametric perturbations can induce accumulation of A*β* fibers: role of clearance processes and astrocytes

The physiological processes, that have been taken into consideration in the model, might encounter a plethora of biological perturbations which may reflect as alteration in the system parameters in a time-dependent manner. Such systemic perturbations can arise from inflammatory conditions or even from a new sleeping habit. Any of our model parameters can get altered for a certain duration due to such a change in the micro-environment of the brain. As the settling time and the steady-state concentration of fA*β* are both dependent on the critical sA*β* production rate *R*_c_, any time-dependent or persistent perturbation in the system parameters resulting in even a slight change of *R*_c_ may have a significant impact on the amount of fA*β*. Whether the system would evolve to a phase dominated by fA*β* and how rapidly the system would recover from the effect of such a transient modification in a system parameter, crucially depend on the change in *R*_c_ due to the parametric perturbation. If *R* exceeds the modified *R*_c_, the fibrillar phase starts to dominate resulting in a rapid production of fA*β*. The recovery time *t*_recov_ which is defined as the time required for the system variable *F* to settle to 99% of its original steady state value (before parametric perturbation), may be relatively longer if the system faces a strong or a long perturbation.

To illuminate this, as an example, we have reduced the microglia density, *M*, to one tenth (1*/*10) of its original value for several perturbation period *t*_pert_ and analyzed how the concentration of fA*β* evolves as a result of this parametric perturbation both during and after *t*_pert_ (Fig. 4). The time of recovery, *t*_recov_, lengthens as we increase *t*_pert_. Furthermore, we note that the concentration of fA*β* at the end of the perturbation period increases considerably with *t*_pert_ as depicted in the left inset of Fig. 4. This suggests that the possibility of developing Alzheimer’s disease (AD) can be much higher if the microglial density drops abruptly even for a year.

**Figure 4:**
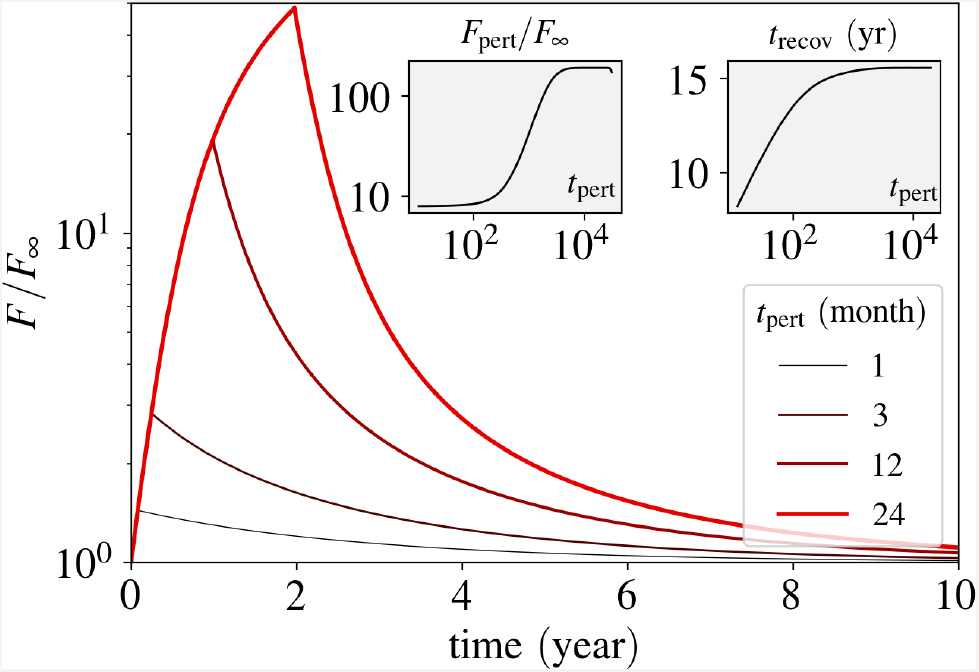
Impact of a perturbation in *M* on fibrillization. During the perturbation time, *t*pert, the number of microglia *M* is reduced ten-fold. *F* is depleted relative to its steady-state value before perturbation. The inset diagram on the left side shows the concentration of fA *β* at the end of the perturbation period, *F*pert. The inset diagram on the right side shows the recovery time, *t*recov, as a function of the perturbation period. The perturbation time in the inset diagrams is plotted in the units of days. *R* = 9 *×* 10^*−*4^.

### Disturbed circadian rhythm can be a key factor in the emergence of AD

As the production and clearance of A*β* both are regulated by the circadian rhythm, it is crucial to consider the impact of a disrupted sleep-wake cycle on A*β* accumulation.

While the model omits an explicit consideration of circadian rhythm in the production rate, it still allows to gain insight into a more realistic setting which includes a circadian rhythm of A*β* production. Moreover, a jejune idea of taking the average of the sA*β* production rate in order to predict the final phase of the system (soluble or fibrillar) does not work in cases where the sA*β* production rate oscillates daily around *R*_c_ due to variations of neural activity in wake and sleep states (Fig. 5-A). Under these circumstances, even a small alteration in the sA*β* production rate may result in a significant change in the settling time close to *R*_c_ thereby making the final phase of the system closely dependent on the sleep-wake cycle.

**Figure 5:**
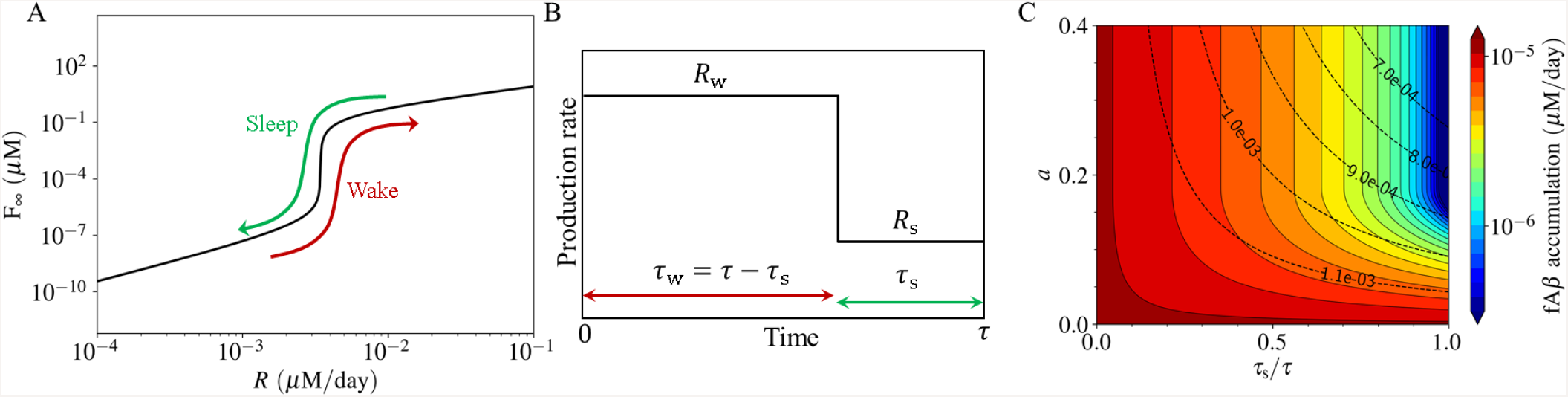
Impact of the circadian rhythm on the fibrillization process. (A) During the sleep-wake cycle, the system can pass through the critical production rate. In such cases, the circadian cycle of A *β* production alleviates a persistent fibrillization. (B) The production rate shown in the course of the circadian time period, *τ* (*∼* a day). It is represented by a periodic rectangular function, with the wake production rate *R*_w_ and the sleep production rate *R*_s_ during the wake and sleep periods, *τ*_w_ and *τ*_s_. (C) The estimated daily fA *β* accumulation as a function of the normalized circadian amplitude *a* = (*R*_w_ *− R*_s_)*/*(*R*_w_ + *R*_s_) and sleep period *τ*_s_, for the case *R*_w_ = 1.2 *×* 10^*−*3^ (*μ*M*/*day). The dashed contour lines show the mean daily production rate *R* = (*R*_w_ *τ*_w_ + *R*_s_*τ*_s_)*/τ* in units of *μ*M *·* day^*−*1^.

This regulatory aspect is essential as the production rate of sA*β* is estimated to lie in a physiological plausible range (the red-highlighted vertical bands in Fig. 2) which includes its critical value *R*_c_. We assume that the diurnal rhythm of overall neural activity results in a diurnal pattern in the production rate of sA*β*, which is represented by a rectangular periodic function 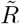 for mathematical simplicity (Fig. 5-B). 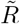 fluctuates between the wake (w) and sleep (s) courses, each with a specific sA*β* production rate *R*_w_ and *R*_s_, and a duration of *τ*_w_ and *τ*_s_, respectively. We define the normalized circadian amplitude as *a* = (*R*_w_ − *R*_s_)*/*(*R*_w_ + *R*_s_). Since both the settling time and the steady-state value of fA*β* concentration depend on the production rate sA*β*, the mean accumulation of fA*β* is determined by *a* and *τ*_s_, especially when the production rate is in a close neighborhood of *R*_c_. The daily accumulation of fA*β* is estimated as 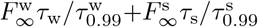, and depicted in Fig. 5-C. The settling time to reach the steady-state value *F_∞_*, varies widely for different 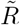 around the critical point *R*_c_. As can be seen from Fig. 5-C, accumulation of fA*β* per day is significantly reduced by having proper relaxation of the overall neural activity, for instance through sleeping. Although the circadian amplitude *a* also plays a pivotal role in this regard, *τ*_s_ outweighs the impact of *a* as long as *a* is high enough to put the production rate of sA*β* below *R*_c_ during sleep. Thus, a good sleep hygiene can be an important protective factor to curtail the accumulation of A*β* in its fibrillar form, thereby averting the onset of AD.

### Potential intervention strategies to prevent or delay fibrillization by elevating *R*_c_

We next consider several potential intervention schemes, reducing fA*β* accumulation in the brain to prevent/treat AD by elevating the critical sA*β* production rate *R*_c_. In a neuro-pathological milieu, different drugs may affect different parameters of the model. An impact of a drug on the parameter *x* can be modeled as either (1 −*ε*_x_)*x* for a drug that decreases the value of a parameter (inhibitory drug), where *ε*_x_ ∈ [0, 1] or (1 + *ε*_x_)*x* for a drug that enhances the value of a parameter (stimulatory drug), where *ε*_x_ *≥* 0, *ε*_x_ being the efficacy of the drug on parameter *x*. Hence, in order to lift *R*_c_, the potential strategies should have an inhibitory effect for the parameters *R* and *k*_p_ (e.g. *R* → (1 − *ε*_R_)*R*), while the rest of the system parameters are to be under a stimulatory treatment (e.g. *M →* (1 + *ε_M_*)*M*).

First, we consider a “monotherapy” that alters only one parameter in the system. Defining *R*_cn_ as the nominal value of *R*_c_ without treatment, Fig. 6-A shows the variation of *R*_c_*/R*_cn_ for several treatment methods corresponding to the efficacy parameters in the range *ε*_x_ *∈* [0, 1]. As can be seen from Fig. 6-A, a reduction in *R* or *k*_p_ has the strongest impact on *R*_c_ because *R*_c_ *≈ R*_cn_(1 *− ε*_x_)^*−*1^ for both *x* = *R* and *x* = *k*_p_.

**Figure 6:**
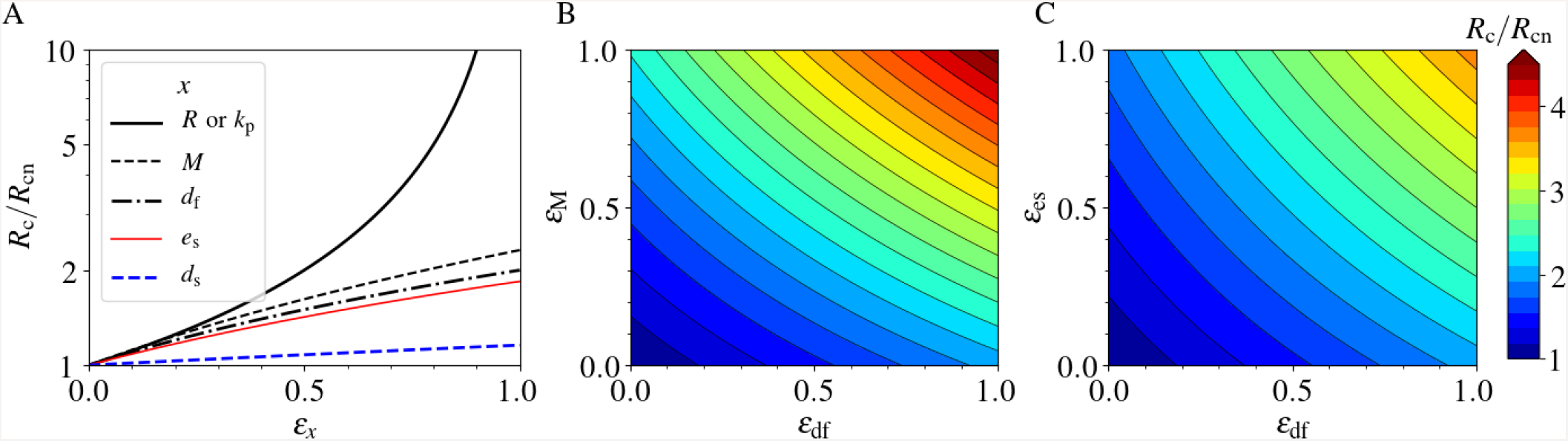
Intervention strategies. The critical production rate *R*_c_, normalized to its nominal value *R*_cn_, as a function of drug efficacies. The treatment strategy is increasing the value of *R*_c_ as much as it exceeds the actual production rate *R*. This happens when a drug decreases the parameters *R* or *k*_p_, or increases any other parameter. *ε*x is the efficacy of a drug, applying to parameter *x*. (A) *R*_c_*/R*_cn_ for single therapies. (B) and (C) *R*_c_*/R*_cn_ for two different combination therapies.

The simultaneous incorporation of two or more such schemes in a “combination therapy” may significantly enhance *R*_c_. We note that the impact emanated from inhibition of *R* and that of *k*_p_ on *R*_c_ are too strong to be achieved with the “combination therapy” with any two of the other parameters. Fig. 6-B and -C show changes of *R*_c_ due to stimulatory effect on the parameter pairs *d*_f_ -*M*, and *d*_f_ -*e*_s_, respectively. As expected, the “combination therapy” averts AD by increasing *R*_c_ remarkably in a nonlinear manner. For instance, where a stimulatory “monotherapy” either on *M* or *d*_f_ with *ε_x_* = 1 hardly increases *R*_c_ by 2-fold, a combination of both of these results in around 4.5-fold enhancement of *R*_c_ (see Fig. 6-B) pointing towards the efficacy of such an approach. Moreover, a “combination therapy” on two parameters which do not play a significant role apiece, may have higher impact than a “monotherapy” on a more influential parameter. For example, a combination of stimulatory therapy on *d*_f_ and *e*_s_ compensates their individual lower impact on increasing *R*_c_ as compared to the “monotherapy” on *M* (compare Fig. 6-C and -A). Our simple model, thus, offers key insight into several options to prevent and treat neuro-inflammatory conditions in the context of fA*β* accumulation in the brain.

## Conclusion

We have developed a minimal ODE-based model of amyloid *β* (A*β*) fibrillization in order to understand the initiation of Alzheimer’s disease (AD), based on the amyloid cascade hypothesis [45, 2, 57]. Our theoretical framework is based on the balance between A*β* production by neurons and their clearance by microglia or physical discharge through the cerebrospinal fluid (CSF). The model suggests that an imbalance between the production and clearance of A*β* may lead to the formation of fibrillar amyloid plaques (fA*β*) which sediment in the brain. Once formed, fA*β* are not only more persistent against clearance by microglia, but also trigger the close-by soluble A*β* oligomers (sA*β*) for a fast fibrillization. Therefore, the homeostatic state of the brain shifts toward the dominance of fA*β* over sA*β*, a process associated with the development of Alzheimer’s disease (AD) in a long run.

Our model predicts two distinct phases for the system, one dominated by sA*β* oligomers and the other dominated by fA*β* plaques. It is argued that if the system finds itself in the latter phase, the amyloid plaques gradually form leading to the onset of AD. The two phases are separated by a critical A*β* production rate, *R*_c_. The phase change occurs when the production rate *R* is greater than *R*_c_. If the physiological value of *R* is very close to *R*_c_, a perturbation in the production or the clearance of A*β* may trigger the phase change in the system and initiate fibrillization. A retrieve from the fibrillar phase might take years according to our results, implying that the system may not be able to find back its original “healthy” homeostatic state.

The production of A*β* is related to the synaptic activity of neurons [16, 17]. We have studied how the lack in relaxation of the neural activity facilitates the onset of A*β* fibrillization. Various physiological conditions, such as neuroinflammation, trauma, or even disturbances in sleep may affect the neural activity and indirectly lead to the accummulation of A*β* [10]. For instance, a flu infection has been recently shown to have an influence on the activation of microglia and neural impairment on mice long after disease recovery [58], indicating that an acute infection may lead to a long-term neuroinflammation which in turn changes the brain homeostatic state and neural activity. Our results suggest that short-term perturbations in the system parameters may result in an abrupt increase in A*β* levels which can persist for years after perturbation. Specifically, we used our model to reveal how sleep might play a key role in preventing the development of AD, by reducing the production of A*β* oligomers. These results are in accordance with the observation that AD patients had sleep disturbance problems years before they show symptoms of AD [9, 10].

Since a reduction of *R*_c_ could initiate progressive fibrillization of A*β*, the onset of AD could be hindered by the treatment strategies that increase the value of *R*_c_. Such an analysis guides us toward new therapeutic or prevention strategies. Efflux of sA*β* through CSF and the clearance by microglia and astrocytes are the primary ways by which the system can get rid of A*β*. Both of these ways could be modulated by drugs, diseases, and injuries. For example, the efficiency of microglia in fA*β* clearance through phagocytosis may be regulated by drugs [59]. These effects could be translated into higher fA*β* decay rate. However, one treatment strategy could be more effective than another if the system has a stronger response to it, which is described by the scaling of *R*_c_ with respect to different parameters. The model shows that *R*_c_ scales superlinearly with the number of microglia, linearly with their efficiency in clearing A*β* for either the soluble or fibrillar form, and linearly with the efflux rate of sA*β* in CSF. It implies that triggering microglial proliferation is more effective in increasing *R*_c_ than stimulating their efficiency in phagocytosis.

According to the model, the direct regulation of sA*β* production or polymerization rate of fA*β* would be very effective in the prevention of fibrillization and AD onset. First, the impact of Apolipoprotein E4 allele (ApoE4) on the disease progression is as important as the production rate of A*β* because ApoE4 directly affects the polymerization rate [60, 61, 62]. It implies that blocking ApoE4 protein should be one of the most influential intervention strategies for the individuals which carry it. Second, since further reduction of the polymerization rate may require the recruitment of astrocytes or changing the chemical properties of the brain parenchyma, the side effects of such a therapy might overweigh its benefit. As changing *R*_c_ is difficult, the regulation of A*β* production seems to be the best solution with the least side effects on the system. The production of A*β* is directly related to the expression of synaptic amyloid precursor protein (APP) which gives rise to A*β* if cleaved abnormally [63]. In addition, APP cleavage products alter synaptic plasticity and activity [64, 65], meaning that A*β* production is not only affected by neural activity, but also affects itself. If such a relation is known, the production of A*β* can be regulated by neural stimulation which brings about broad new therapeutic schemes focusing on electrical stimulation of neurons to regulate A*β* production for AD prevention.

All in all, as Karran et al. reviewed in 2011 [66], the complexity of the disease has not been understood completely such that many clinical trials focusing on reducing the production of A*β* failed because the disease had already shifted to a new phase, where many other pathological and neuroinflammatory factors entered. In this regard, it is difficult to directly relate the model results to clinical trials. Nevertheless, the model predicts which treatment strategies could be more effective, essentially before the onset of the disease. For instance, we can safely argue that targeting less amyloid production at the first place is equally effective as avoiding ApoE4, but far more beneficial than the clearance through antibodies due to the scaling laws we have obtained from the model.

## Author Contributions

M.H. and M.M.H. designed the research. M.H. developed the mathematical model. M.H., S.K., G.M., and T.M. did the analysis and investigation. M.M.H. supervised the research. All authors contributed in discussions and wrote the manuscript.

## Acknowledgments

We express our gratitude to the authors of Ref. [30] and Dr. Georg Meisl for providing their experimental data. M.H, S.K., and M.M.H. acknowledge the support by the Helmholtz Association, Zukunftsthema “Immunology and Inflammation” (ZT-0027). G.M. and M.M.H. are also thankful for the support by German Federal Ministry of Education and Research within the measures for the establishment of systems medicine, eMed project SYSIMIT (FKZ.01ZX1608B).

## Supplementary Materials

Supplementary Materials accompany this paper.

