## Supplementary Materials for "Mathematical model shows how sleep may affect amyloid *β* fibrillization"

### Supplementary Materials for "Mathematical model shows how sleep may affect amyloid $\beta$ fibrillization"


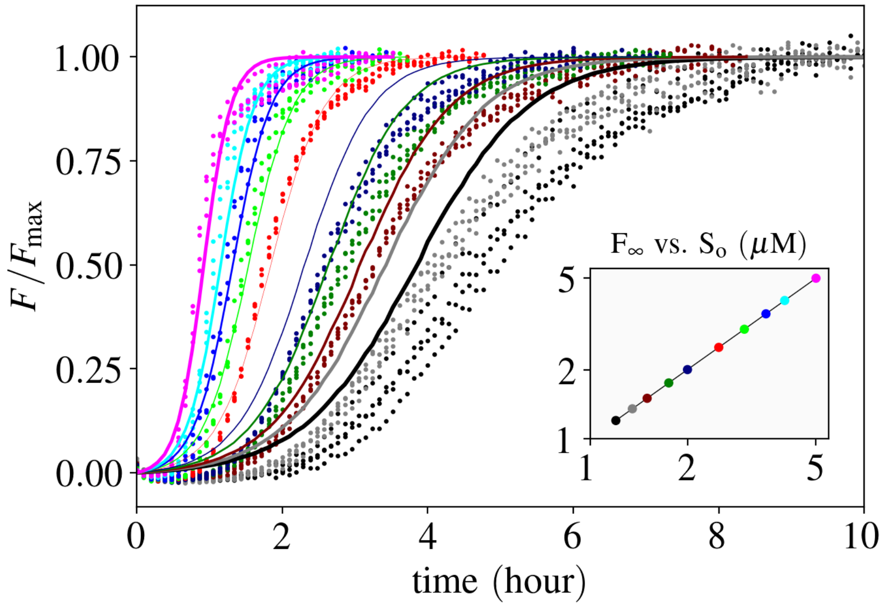


**Figure S1: Estimation of the nucleation and polymerization rates according to the ThT experiments conducted by Cohen et al. 2013 [30]. The points and lines correspond to the experimental data and the model, respectively. The colors in the main plot correspond to the marker colors in the inset. A reduced model is used for the estimation, where only an initial concentration for sA$\beta$ ($S_\mathrm{o}$) and the nucleation $k_\mathrm{n}$ and polymerization rate constant $k_\mathrm{p}$ are taken into account, i.e. Eq. 1 and 2 reduce to $\dot{S} = -k_\mathrm{p}SF - k_\mathrm{n}S^2$ and $F = S_\mathrm{o} - S$. Here, $S \equiv [\mathrm{sA}\beta]$. The parameters are estimated through the differential evolution algorithm [37] with logarithmic weight $\log S_\mathrm{o}$ on the cost function. $k_\mathrm{n} = 1.7 \times 10^{-1}\ \mu\mathrm{M}^{-1} \cdot \mathrm{day}^{-1}$ and $k_\mathrm{p} = 2.6 \times 10^{1}\ \mu\mathrm{M}^{-1} \cdot \mathrm{day}^{-1}$. Our proposed simple model is able to catch the fibrillization behavior qualitatively.**

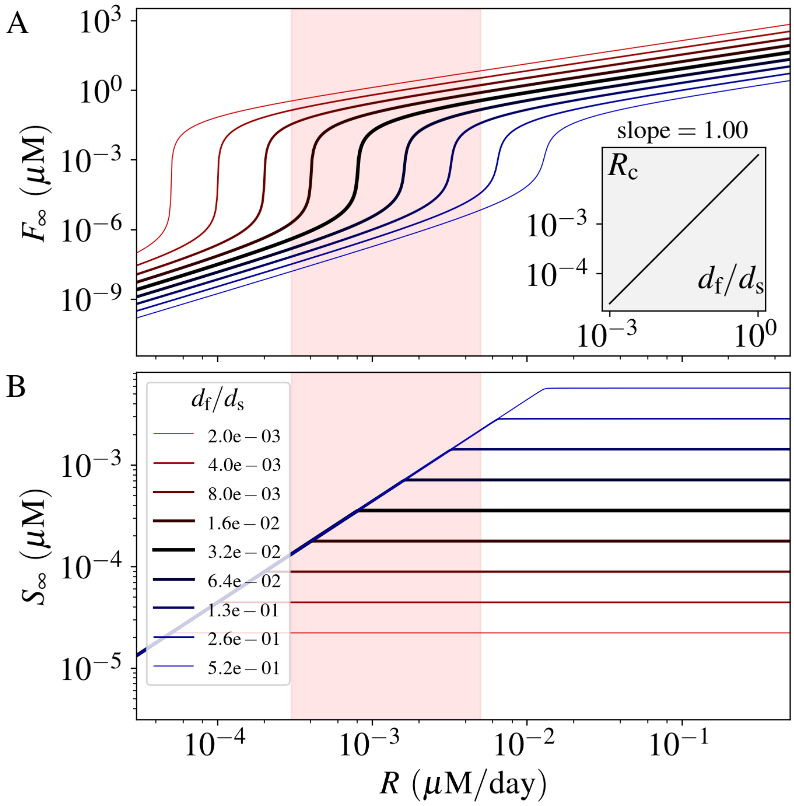


**Figure S2: Impact of the fA$\beta$ decay rate $d_f$ on the fibrillization of A$\beta$. Panel A and B show the concentration of fA$\beta$ and sA$\beta$, respectively. The inset diagram shows that $R_c$ scales linearly with the changes of $d_f$. $d_s$ is only used for the normalization purpose and kept constant.**

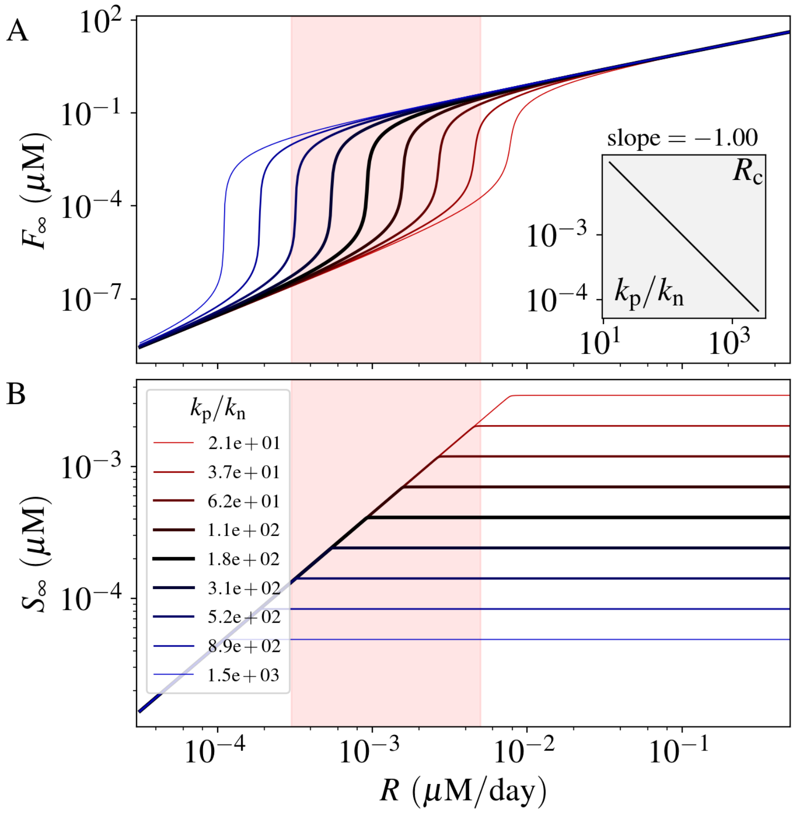


**Figure S3: Impact of the polymerization rate $k_\mathrm{p}$ on the fibrillization of A$\beta$. Panel A and B show the concentration of fA$\beta$ and sA$\beta$, respectively. The inset diagram shows that $R_\mathrm{c}$ scales inversely with $k_\mathrm{p}$. $k_\mathrm{n}$ is only used for the normalization purpose and kept constant. Reduction of $k_\mathrm{p}$ is related to the number and efficiency of astrocytes.**

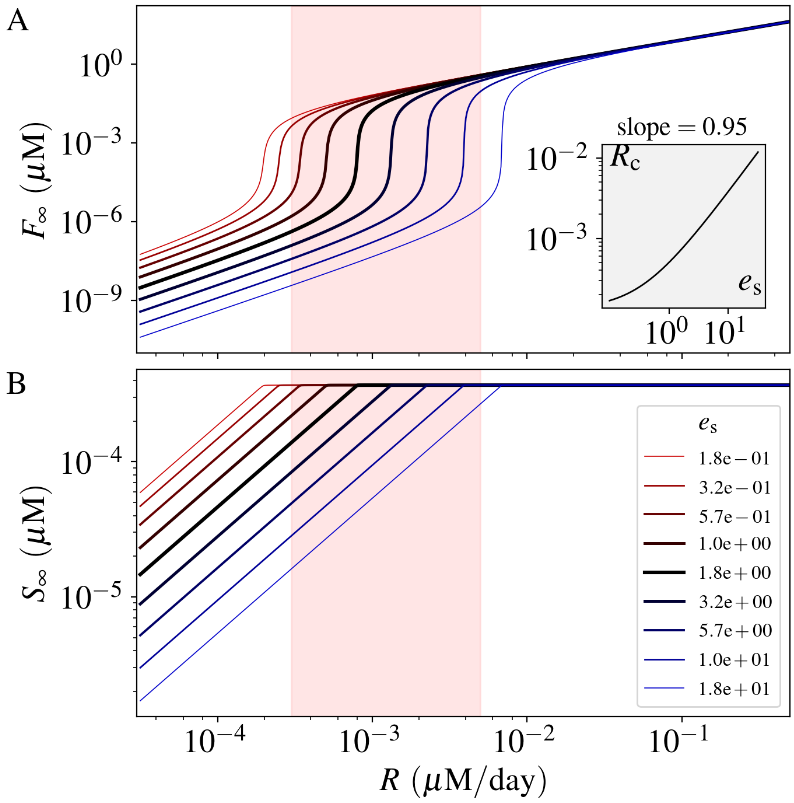


**Figure S4: Impact of the sA$\beta$ efflux rate $e_s$ on the fibrillization of A$\beta$. Panel A and B show the concentration of fA$\beta$ and sA$\beta$, respectively. The inset diagram shows that $R_c$ scales almost linearly with $e_s$.**
